# The MCMV immunoevasin gp40/*m152* inhibits NKG2D receptor RAE-1γ by intracellular retention and cell surface masking

**DOI:** 10.1101/2020.11.17.386763

**Authors:** Natalia Lis, Zeynep Hein, Swapnil S. Ghanwat, Venkat Raman Ramnarayan, Benedict J. Chambers, Sebastian Springer

## Abstract

NKG2D is a crucial Natural Killer (NK) cell activating receptor, and the murine cytomegalovirus (MCMV) employs multiple immunoevasins in order to avoid NKG2D-mediated activation. One of the MCMV immunoevasins, gp40 (*m152*), downregulates the cell surface NKG2D ligand, RAE-1γ, thus limiting NK cell activation. This study establishes the molecular mechanism by which gp40 retains RAE-1γ in the secretory pathway. Using flow cytometry and pulse chase analysis, we demonstrate that gp40 retains RAE-1γ in the early secretory pathway, and that this effect depends on the binding of gp40 to a host protein, TMED10, a member of the p24 protein family. We also show that the TMED10-based retention mechanism can be saturated, and that gp40 has a backup mechanism as it masks RAE-1γ on the cell surface, blocking the interaction with the NKG2D receptor and thus NK cell activation.

**Summary statement:** MCMV immunoevasin gp40 inhibits the NKG2D-activating ligand RAE-1γ by intracellular retention that depends on the p24 member TMED10, and additionally by masking it at the cell surface.

## Introduction

Human Cytomegalovirus (HCMV)^1^ infections are widespread among humans, with a seroprevalence of about 50% (or higher in old persons and in developing countries). Though mostly asymptomatic, HCMV infection is a serious threat to immunocompromised patients. It can develop into a serious disease that damages the nervous system, retinal cells, gastrointestinal tract, and lungs (Dioverti and Razonable, 2016).

HCMV is strictly species-specific. Therefore, to study the infection and immune evasion strategies, it is common to use the similar murine cytomegalovirus (MCMV). The MCMV model allows to study the impact of viral factors on the host *in vivo*, and discoveries made with MCMV have often generated hypotheses for HCMV infection (Brizić et al., 2018; Hummel and Abecassis, 2002; Reddehase and Lemmermann, 2018).

CMV expresses a large number of immunoevasins, i.e., viral proteins that help the virus to evade the host immune response. Most notably, some CMV immunoevasins have evolved to hamper both lines of the host defense, the innate and the adaptive immune responses. One such example is the MCMV glycoprotein gp40 (encoded by *m152*). gp40 is expressed starting about three hours after infection, and it antagonizes three types of molecules: first, T-cell activating MHC class I proteins, second, the protein stimulator of interferon genes (STING), and third, the natural killer (NK) cell-activating stress marker, RAE-1 (Lodoen et al., 2003; Stempel et al., 2019; Ziegler et al., 1997). It is not clear whether gp40 targets all these molecules simultaneously or at different times post-infection, but learning about molecular details of gp40 action will lead to a better understanding of viral adaptation and co-evolution with the host.

RAE-1 is a GPI-anchored protein, and it has five isoforms (α, β, γ, δ, ε) that are about 90% similar in the amino acid sequence. It is a stress ligand, a molecule that is not expressed in healthy cells of adults, but its expression can be triggered by viral infection or tumorigenesis (Cerwenka and Lanier, 2001; Lanier, 2015). Once present on the cell surface, RAE-1 binds to the NKG2D receptor, which is one of the main activating NK cell receptors, and plays a central role in the defense against viruses (Chan et al., 2014; Eagle and Trowsdale, 2007; Raulet, 2003).

The significance of RAE-1/NKG2D interaction in the context of MCMV infection has been previously recognised. During MCMV infection, gp40 downregulates cell surface RAE-1 and retains it in the early secretory pathway, resulting in reduced NK cell activation and higher viral titers. Both proteins bind to each other *in vitro*, with some RAE-1 isoforms being more susceptible to gp40 than others (Arapović et al., 2009b; Zhi et al., 2010); but the mode of interaction between gp40 and RAE-1 in cells, as well as the molecular mechanism of gp40-driven downregulation of cell surface RAE-1, remain unknown.

Here, we investigate RAE-1γ, the isoform of RAE-1 that is most susceptible to gp40 downregulation, and we identify a previously unknown molecular mechanism of RAE-1γ retention, which is bridging RAE-1γ to the host protein carrying retention/retrieval motifs, TMED10. We also show for the first time a second mechanism of inhibiting NK cell activation by NKG2D recognition, namely the masking of cell surface RAE-1γ by gp40.

## Results

### MCMV gp40 downregulates RAE-1γ cell surface levels and retains it in the early secretory pathway

The NK cell-activating ligand RAE-1 is not or only weakly expressed in normal cells, but it is induced by stress and viral infection (Cerwenka et al., 2000; Lodoen et al., 2003). In order to study the influence of gp40 on RAE-1γ, we established cell lines that express N-terminally HA-tagged RAE-1γ (HA-RAE-1γ) alone or together with gp40. We wished to test the influence of gp40 on RAE-1γ in the absence or presence of MHC class I molecules (another gp40 target), and therefore we chose three different combinations of genes and cells: first, K41 murine fibroblasts expressing HA-RAE-1γ and gp40; second, B78H1 murine melanoma cells (lacking MHC class I) expressing HA-RAE-1γ and gp40; and third, HEK293T human cells (with human MHC class I molecules that are not affected by gp40) expressing with HA-RAE-1γ and with gp40 in a vector that expressed GFP from an internal ribosomal entry site (IRES) (Table S1).

To test the impact of gp40 on the cell surface level of RAE-1γ, we stained these cells for cell surface RAE-1γ and analyzed them by flow cytometry using the monoclonal anti-RAE-1γ antibody, CX1. For all flow cytometry data, we used cells with transfected with empty vectors as controls, and we normalized our results to those obtained for cells expressing HA-RAE-1γ without gp40 (Original data Table S2).

In the cells expressing gp40, we found that HA-RAE-1γ cell surface levels were reduced (Fig. 1A). As observed before, in K41 cells, gp40 downregulated also endogenous MHC class I molecules (Fig. S1A) (Janßen et al., 2016). Additionally, K41 cells expressed small amounts of endogenous RAE-1γ, which was also downregulated by gp40 (Fig. S1B). In murine and human cell lines, HA-RAE-1γ expression was reduced (Fig. 1B).

**Figure 1.**
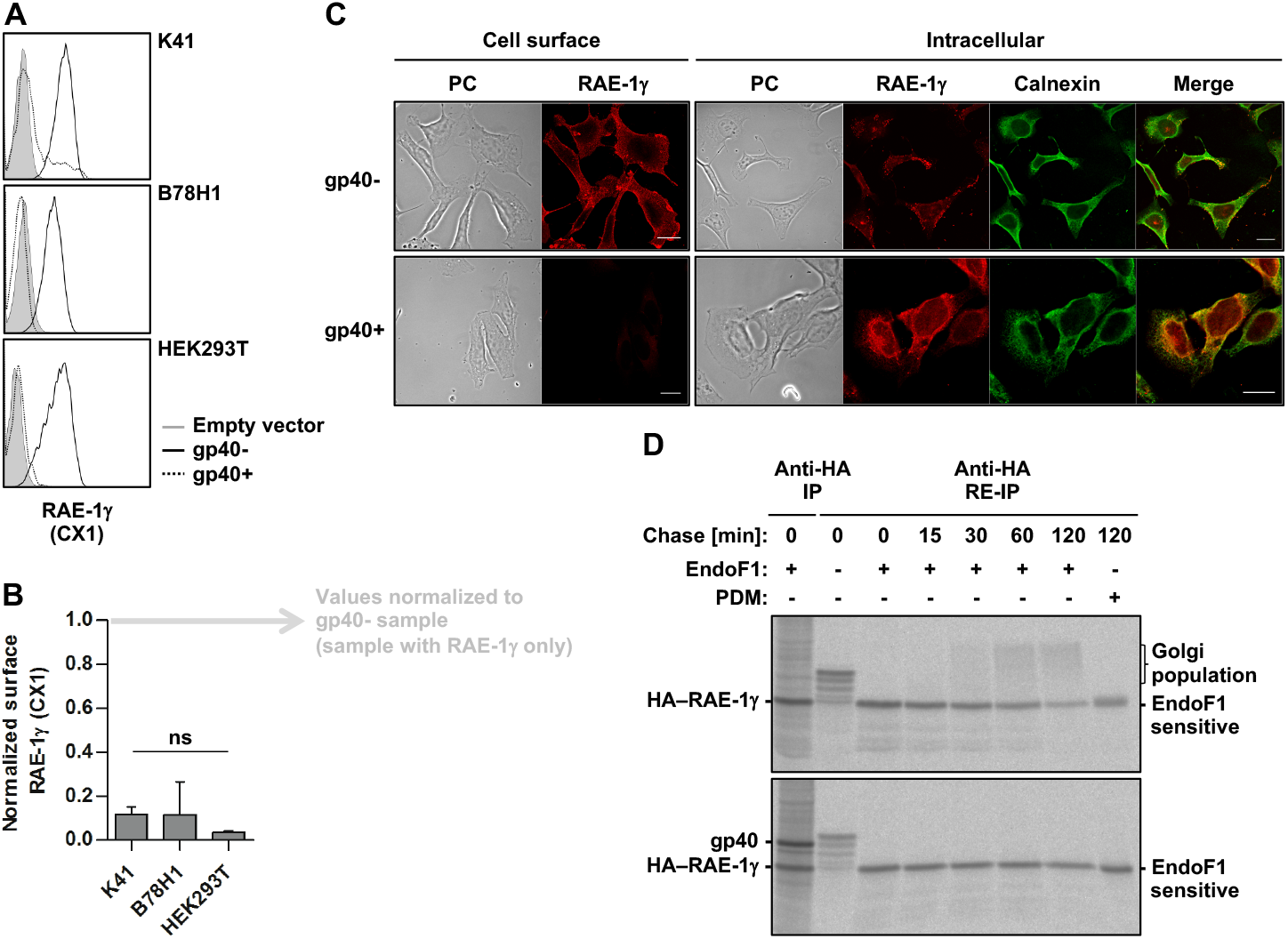
MCMV gp40 downregulates RAE-1γ cell surface level and retains it in the early secretory pathway. A-B) K41, B78H1 and HEK293T cells were transfected with empty vector or HA-RAE-1γ alone (gp40-), or together with gp40 (gp40+), and the surface expression of HA-RAE-1γ was determined by staining with CX1 and flow cytometry. Grey shading: cells expressing empty vector (B78H1 and HEK293T cells) or antibody control (K41 cells). One representative experiment out of three is shown. The mean fluorescence intensity of cell surface HA-RAE-1γ represented in a bar chart (B), where the values were normalized to the HA-RAE-1γ mean fluorescence intensity in the gp40-cells (mean ± SD, n=3, ns P > 0.05). C) B78H1 cells transfected with HA-RAE-1γ alone (gp40-), or together with gp40 (gp40+), were fixed, permeabilized (intracellular stain), and stained with anti-HA antibody followed by mAlexa488, or anti-calnexin antibody, followed by rCy3. Scale bar, 20 μm. D) HEK293T cells were transfected with HA-RAE-1γ alone, or with gp40, were pulse-labeled for 10 min and chased for the indicated times. Cells were lysed in 1% Triton X-100 buffer, and the proteins were immunoprecipitated from the lysate with an anti-HA antibody. Immunoprecipitates (excluding the first lane) were dissociated with denaturation buffer, and proteins were re-immunoprecipitated with an anti-HA antibody (RE-IP). Samples were digested with EndoF1 or Protein Deglycosylation Mix, which removes all N-linked and O-linked glycosylation (PDM), and separated by SDS-PAGE.

To identify the intracellular steady-state localization of HA-RAE-1γ, we performed immunofluorescence microscopy in B78H1 cells using an anti-HA antibody. Without gp40, HA-RAE-1γ was detectable at the cell surface and to some extent in vesicular structures near the plasma membrane, and it did not co-localize with the ER marker. In cells expressing gp40, HA-RAE-1γ was no longer visible at the cell surface but instead accumulated in the ER (Fig. 1C).

In order to understand the dynamics of RAE-1γ transport throughout the early secretory pathway, we then performed pulse-chase and immunoprecipitation analysis combined with endoglycosidase F1 (EndoF1) digestion. Like Endoglycosidase H (EndoH), EndoF1 removes glycans from proteins that are located in the pre-medial Golgi compartments (Trimble and Tarentino, 1991). Due to the action of mannosidase II, most glycans become resistant to EndoF1 in the medial Golgi. In the *trans*-Golgi, glycans are further modified by the addition of sialic acid residues (Fritzsche and Springer, 2013). Thus, the intracellular location of a protein determines its glycosylation, which can be easily visualized as size shift in SDS-PAGE.

We radiolabelled HEK293T cells expressing HA-RAE-1γ and gp40 and chased for up to two hours. We lysed the cells in 1% Triton X-100 and performed immunoprecipitations with an anti-HA antibody. Since the complex of RAE-1γ and gp40 was resistant to 1% Triton X-100 (Fig. 1D, first lane of lower panel), we dissociated it by boiling the immunoprecipitates in 2% SDS and then re-immunoprecipitated RAE-1γ via its HA tag.

In cells without gp40, HA-RAE-1γ matured rapidly, as shown by the decrease in the EndoF1-sensitive signals as early as 15 minutes into the chase. After 30 minutes, we observed a diffuse high molecular weight band that corresponds to the sialylated *trans*-Golgi population of HA-RAE-1γ (Fig. 1D, top panel) (Fritzsche and Springer, 2013). Within 120 minutes, 65% of the protein became EndoF1-resistant. The sample in the last lane represents the total protein amount after 120 minutes irrespective of its cellular location and glycosylation state. The quantification suggests that about 50% of HA-RAE-1γ was degraded within two hours (Fig. S1C, samples labeled #). In contrast, in the cells expressing gp40, the levels of EndoF1-sensitive HA-RAE-1γ remained nearly constant throughout the chase, and the diffuse *trans*-Golgi sialylated signal was absent even after 120 minutes (Fig. 1D, bottom panel). HA-RAE-1γ was also more stable, and at the end of the chase, only about 20% of the protein was degraded (Fig. S1C #). The results confirm, in line with previous reports, that gp40 decreases the cell surface expression of RAE-1γ and retains it in the early secretory pathway (Arapović et al., 2009b; Lodoen et al., 2003).

### RAE-1γ retention depends on the gp40/TMED10 interaction

We recently established the molecular mechanism of gp40-driven MHC class I retention (Janßen et al., 2016; Ramnarayan et al., 2018). To maintain its own localization in the early secretory pathway, gp40 binds to the host proteins of the p24 family that are involved in vesicular ER-Golgi protein trafficking. The interaction depends on the linker that connects the luminal and the transmembrane domains of gp40, and it is the strongest between gp40 and the transmembrane p24 trafficking protein 10 (TMED10) (Ramnarayan et al., 2018). In contrast, interaction between gp40 and STING does not depend on the gp40 linker and presumably binding to TMED10 or other p24 proteins (Stempel et al., 2019).

We therefore asked whether gp40 uses its interaction with TMED10 to retain RAE-1γ. To test whether TMED10 is present in the complex with RAE-1γ and gp40, we transfected wild type HEK293T cells with HA–RAE-1γ alone or together with gp40, lysed cells in a mild detergent (digitonin), and performed co-immunoprecipitation using an anti-HA antibody. Immunoblot analysis showed that TMED10 is co-immunoprecipitated with RAE-1γ in cells that express gp40 (Fig. 2A). In the absence of gp40, RAE-1γ – just like class I – did not bind to TMED10. This suggests that gp40 ties RAE-1γ to TMED10 in order to retain it in the ER (Ramnarayan et al., 2018).

**Figure 2.**
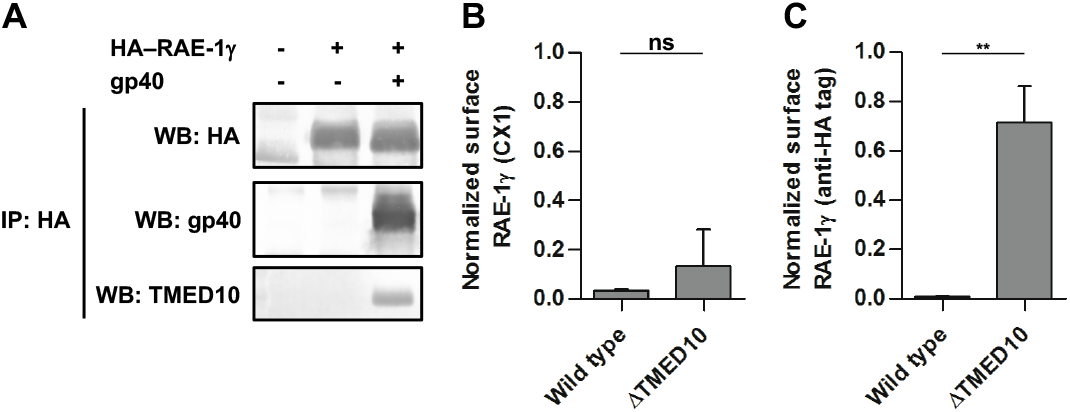
RAE-1γ retention depends on gp40/TMED10 interaction. A) HEK293T cells were transfected with empty vector, or HA-RAE-1γ alone, or together with gp40. Next, cells were lysed in 1% digitonin, and the proteins were immunoprecipitated from the lysate with an anti-HA antibody. Samples were digested with PDM, separated by SDS-PAGE, and immunoblotted against HA tag, gp40, and TMED10. B-C) Wild type HEK293T cells, or HEK293T^Δ^TMED10^^ cells were transfected with empty vector, or HA-RAE-1γ alone, or together with gp40, as indicated, and surface expression of HA-RAE-1γ was determined by staining with (B) CX1, or (C) anti-HA antibody, followed by APC-conjugated secondary antibody (APC), and flow cytometry. The values were normalized to the HA-RAE-1γ mean fluorescence intensity in the gp40-cells (mean ± SD, n=3, ns P > 0.05; ** P ≤ 0.01).

This observation led us to hypothesize that in cells that lack TMED10, RAE-1γ should not be retained by gp40. We used HEK293T cells with a knockout of TMED10 (HEK293T^*ΔTMED10*^) and measured the surface expression of transfected HA-RAE-1γ in the presence or absence of gp40. To our surprise, however, the TMED10 knockout only restored only 10-20% of the previous surface expression of HA-RAE-1γ as measured with the CX1 antibody (Fig. 2B, Fig. S2A). Based on the gp40/MHC class I studies, we had expected a nearly complete rescue of RAE-1γ upon TMED10 knockout (Ramnarayan et al., 2018).

We therefore decided to repeat this experiment with another antibody, and we took advantage of the fact that our RAE-1γ construct has an HA tag on its extracellular N terminus. Thus, we repeated the flow cytometry analysis using the anti-HA antibody. Interestingly, the anti-HA staining in HEK293T^*ΔTMED10*^ cells expressing gp40 showed that about 80% of HA-RAE-1γ was rescued (Fig. 2C, Fig. S2B). This result suggests that when gp40 is expressed in *TMED10*-deleted cells, the majority of the HA-RAE-1γ that is seen on the cell surface with the anti-HA antibody is not recognized by the CX1 antibody. The simplest explanation for this discrepancy is that the majority of the HA-RAE-1γ on the cell surface is masked or altered and thus no longer detectable by the CX1 antibody.

Taken together, we conclude that TMED10 is involved in the retention of RAE-1γ by anchoring the complex of gp40 and RAE-1γ in the ER. Moreover, in the absence of TMED10, we observed an additional and previously unknown adverse effect of gp40 on the cell surface RAE-1γ that precludes recognition of RAE-1γ by the CX1 antibody.

### The gp40 linker mutant as a model to study gp40/RAE-1γ interaction in the absence of the ER anchoring

The TMED10 protein is central for the ER-Golgi transport and cargo selection in the early secretory pathway (Lopez et al., 2019; Pastor-Cantizano et al., 2016). Therefore, to avoid any side effects caused by the knockout of TMED10, we decided to proceed instead with a mutant gp40 protein that was unable to interact with it. We replaced HEK293T^*ΔTMED10*^ cells with wild type HEK293T cells expressing a previously described gp40 linker mutant in which the 43 amino acid long linker is replaced by a (Gly4Ser)9 sequence. This gp40 linker mutant (gp40LM) does not bind TMED10 or other p24 proteins (TMED9, TMED5, TMED2) and is not retained in the early secretory pathway (Ramnarayan et al., 2018).

We anticipated that the effect of gp40LM on RAE-1γ in wild type HEK293T cells should phenocopy the effect of wild type gp40 in HEK293T^*ΔTMED10*^ cells, since in both cases, gp40 cannot be anchored by TMED10. Thus, to study the effect of the gp40LM on RAE-1γ cell surface levels, we transfected wild type HEK293T cells with HA-RAE-1γ alone or together with gp40LM. After staining with either CX1 anybody or anti-HA antibody, we again compared the cell surface RAE-1γ levels with flow cytometry. As expected, in gp40LM-expressing cells, HA-RAE-1γ was restored to the cell surface, but the majority of the HA-RAE-1γ molecules detected with the anti-HA antibody was not recognized by the CX1 antibody (70% vs 15%) (Fig. 3B-C). The same effects of gp40WT and gp40LM on HA-RAE-1γ were seen in K41 and B78H1 cells (Fig. S3A-B).

**Figure 3.**
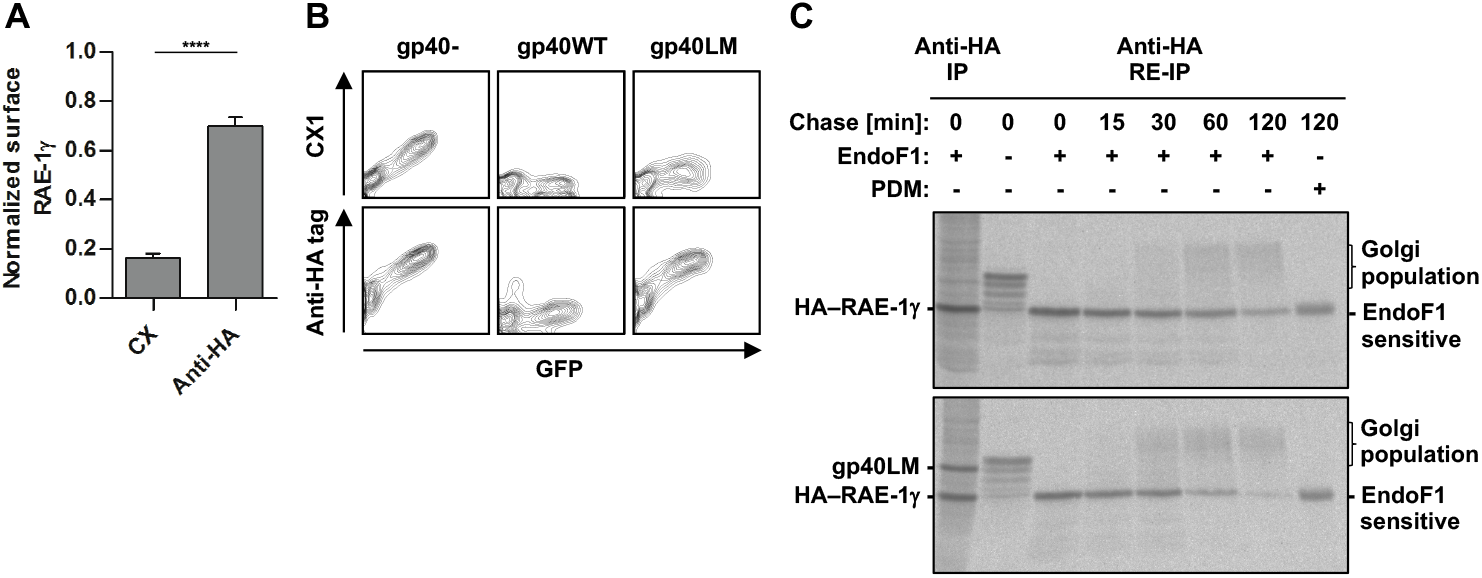
The gp40 linker mutant as a model to study gp40/RAE-1γ interaction in the absence of the ER anchoring. A) HEK293T cells were transfected with empty vector, or HA-RAE-1γ alone, or together with gp40 linker mutant, and surface expression of HA-RAE-1γ was determined by staining with CX1, or anti-HA tag antibody, as indicated, followed by APC, and flow cytometry. The values were normalized to the HA-RAE-1γ mean fluorescence intensity in the gp40-cells (mean ± SD, n=3, **** P ≤ 0.0001). B) One representative experiment out of three is showing HEK293T cells expressing HA-RAE-1γ alone (gp40-), or together with and gp40 wild type (gp40WT) or gp40 linker mutant (gp40LM) in GFP expressing vector. Surface expression of HA-RAE-1γ was determined by staining with CX1, or anti-HA antibody, followed by APC, and flow cytometry. C) HEK293T cells transfected with HA-RAE-1γ alone, or with gp40 linker mutant (gp40LM), were pulsed-labeled for 10 min, and chased for the indicated times. Cells were lysed in 1% Triton X-100 buffer, and the proteins were immunoprecipitated from the lysate with an anti-HA antibody. Immunoprecipitates (excluding the first lane) were dissociated with denaturation buffer, and proteins were re-immunoprecipitated with an anti-HA antibody (RE-IP). Samples were digested with EndoF1 or PDM, and separated by SDS-PAGE.

We also investigated the intracellular trafficking of HA-RAE-1γ in the presence of gp40LM by pulse-chase analysis. HA-RAE-1γ trafficking was identical in cells expressing gp40LM and in cells without gp40. The amount of EndoF1-sensitive RAE-1γ decreased after 15 minutes of chase, and the high molecular weight signals corresponding to the Golgi population appeared after 30 minutes (Fig. 3C, Fig. S3D).

In immunofluorescence microscopy of B78H1 cells, HA-RAE-1γ showed identical localization with or without gp40LM, did not co-localize with an ER marker, and was mostly present at the cell surface (Fig. S3C).

We conclude that just like gp40 in TMED10-deleted cells, gp40LM has no retention effect on RAE-1γ in wild type cells. gp40LM therefore is a suitable model for studying the molecular details of the gp40/RAE-1γ interaction in the absence of TMED10-mediated retention, without the potential side effects on the cell caused by the lack of TMED10. Additionally, this approach eliminates potential interactions between gp40 and any other members of the p24 family that might also function as an anchor for the gp40/RAE-1γ complex (Ramnarayan et al., 2018).

### gp40 and RAE-1γ interact tightly, and both proteins reach the cell surface

Since the complex of gp40 with HA-RAE-1γ is resistant to Triton X-100 lysis (Figure 1D), and thus more stable than the gp40/MHC class I complex, which dissolves in Triton X-100, we next investigated the persistence of the gp40/RAE-1γ complex over time by pulse-chase analysis in wild type HEK293T transfected with HA-RAE-1γ alone or with wild type gp40 (gp40WT) or gp40LM (Janßen et al., 2016; Ramnarayan et al., 2018). Both proteins have a significantly different molecular weight and can thus be followed on one SDS-PAGE gel. gp40 and RAE-1γ bound to each other right after synthesis (zero minutes of chase) and stayed together for at least 120 minutes (Fig. 4A). The gp40WT molecules that co-immunoprecipitated with HA-RAE-1γ acquired partial EndoF1 resistance, suggesting that the complex circulates in the early secretory pathway, which is analogous to our previous observations of the gp40/MHC class I complex (Fig. 4A, middle panel, indicated with arrowheads) (Janßen et al., 2016). We did not observe partial EndoF1 resistance of HA-RAE-1γ, even though it was glycosylated (Fig. S4A). Interestingly, gp40LM was also bound to HA-RAE-1γ throughout the entire chase, suggesting that ER anchoring is not relevant for the complex formation, and that the gp40LM/HA–RAE-1γ complex can reach the cell surface intact (Fig. 4A). The maturation of HA-RAE-1γ itself in the presence of gp40LM was clearly visible (Fig. 3C).

**Figure 4.**
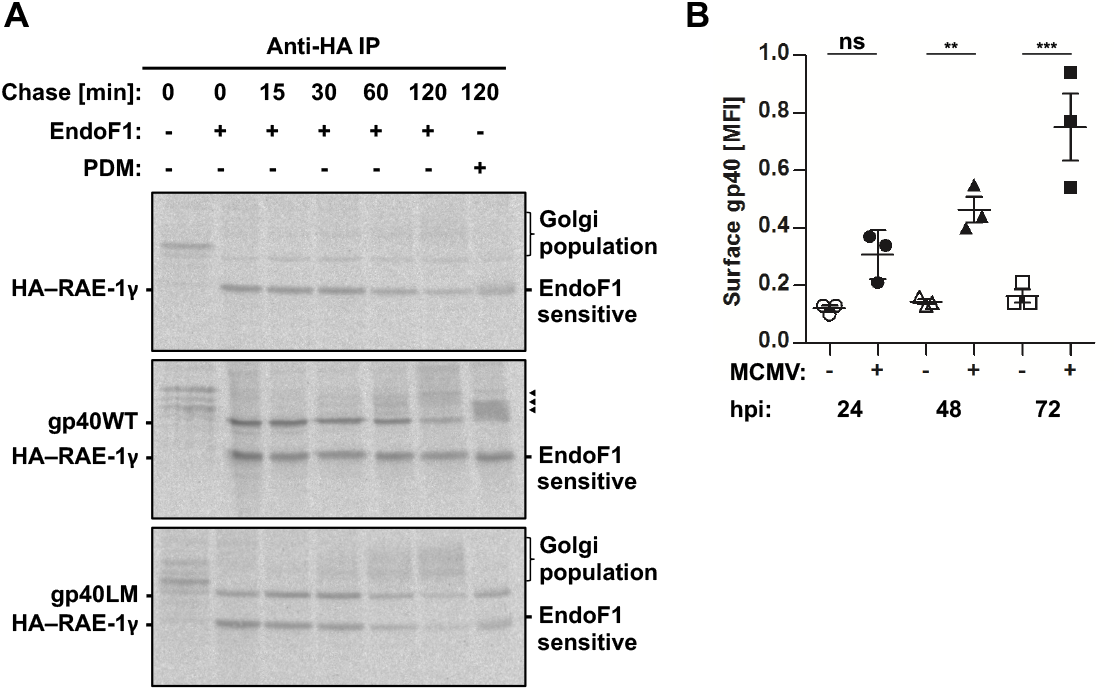
gp40 and RAE-1γ interact tightly and both proteins reach the cell surface. A) HEK293T cells transfected with HA-RAE-1γ alone, or with gp40 wild type (gp40WT) or gp40 linker mutant (gp40LM), were pulsed-labeled for 10 min and chased for the indicated times. Cells were lysed in 1% Triton X-100 buffer, and the proteins were immunoprecipitated from the lysate with an anti-HA antibody. Samples were digested with EndoF1 or PDM and separated by SDS-PAGE. B) K41 cells were infected with MCMV, and incubated for indicated time (hpi, hours post infection). After harvesting, cells were incubated with FcR blocking reagent, and surface expression of gp40 was determined by staining with anti-gp40 antibody, followed by APC, and flow cytometry. The mean fluorescence intensity of cell surface gp40 represented in a scatter plot. The values represent the mean fluorescence intensity (mean ± SD, n=3, ns P > 0.05; * P ≤ 0.05; ** P ≤ 0.01; *** P ≤ 0.001).

The formation of a gp40LM/HA–RAE-1γ complex and its surface transport observed in the pulse chase suggest that some gp40LM should be detectable at the cell surface by antibody staining and flow cytometry. This was indeed the case, and unexpectedly, we also detected gp40WT (Fig. S4B).

All previously published data suggest that gp40 acts in the early secretory pathway in order to antagonize MHC class I molecules, RAE-1, and STING, and upon overexpression is able to reach also the lysosomes (Arapović et al., 2009b; Janßen et al., 2016; Ramnarayan et al., 2018; Stempel et al., 2019; Ziegler et al., 1997). Based on our gp40 surface stains above, we wondered whether during an actual MCMV infection, some cohort of gp40 might fail to be retained and instead proceed to the cell surface. During MCMV infection, viral proteins are often overexpressed, and we speculated that overabundant gp40 might exceed the capacity of cellular p24 proteins to retain it in the ER (Damdindorj et al., 2014). Thus, to test whether gp40 reaches the cell surface during infection, we infected K41 cells with MCMV and stained their surface with an anti-gp40 antibody. To monitor the infection, we stained for the intracellular infection marker, IE1 (Fig. S4C). Indeed, gp40 appeared on the cell surface post-infection, and its amount increased over time (Fig. 4B).

Our data demonstrate that gp40 forms a strong complex with RAE-1γ that circulates in the early secretory pathway. If gp40 cannot bind to its retention factor, TMED10, then the gp40/RAE-1γ complex is still formed, but it is no longer retained in the ER and consequently travels to the cell surface. Some gp40 molecules reach the surface of infected cells, which suggests that escape of the gp40/RAE-1γ complex does indeed occur during MCMV infection.

### gp40 blocks the interaction between NKG2D and RAE-1γ

The crystal structure of the gp40/RAE-1γ complex shows that gp40 binds to the top of the RAE-1γ molecule, forming two interaction surfaces (Fig. 5A) (Humphrey et al., 1996; Wang et al., 2012). Our results show that the complex of gp40 with RAE-1γ indeed reaches the cell surface (Fig. 4), and thus, the simplest explanation for the lack of detection of most of the HA-positive RAE-1γ surface molecules with CX1 is that gp40 masks the CX1 epitope (Fig. 2B-C).

**Figure 5.**
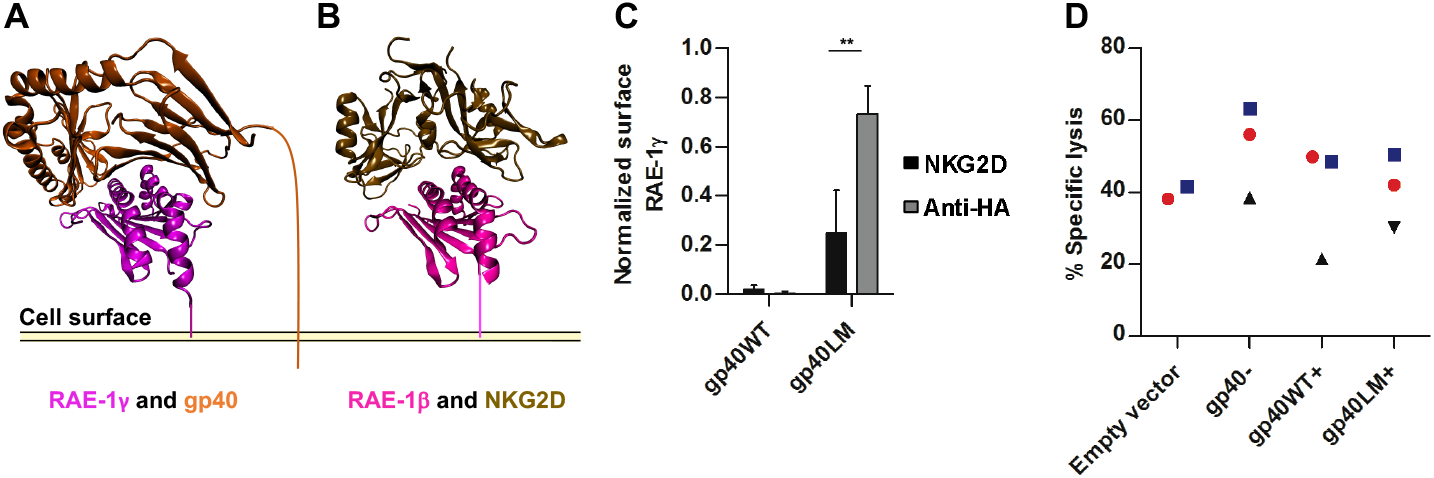
gp40 blocks interaction between NKG2D and RAE-1γ. A) Crystal structure of RAE-1γ in complex with gp40 (PDB 4G59). Membrane anchoring added as schematic. B) Crystal structure of RAE-1β in complex with NKG2D (PDB 4PP8). Membrane anchoring added as schematic. C) HEK293T cells were transfected with HA-RAE-1γ alone (gp40-), or together with gp40 wild type (gp40WT) or gp40 linker mutant (gp40LM), and surface expression of HA-RAE-1γ was determined by staining with anti-HA antibody or with recombinant mouse NKG2D-Fc chimera protein, followed by APC, and flow cytometry. The values were calculated by subtracting the background mean fluorescence intensity and normalized to the HA-RAE-1γ mean fluorescence intensity in the gp40-cells (mean ± SD, n=4, ** P ≤ 0.01). D) B78H1 cells transfected with an empty vector, or HA-RAE-1γ alone (gp40-), or together with gp40 wild type (gp40WT), or gp40 linker mutant (gp40LM) were labelled for one hour with ^51^ Cr, mixed with freshly isolated and pre-activated mouse NK cells at 10:1 ratio, co-incubated for four hours, and the specific cell lysis was calculated (n=2-3).

The gp40/RAE-1γ complex resembles the complex between RAE-1β and the NKG2D receptor homodimer (Fig. 5B)(Humphrey et al., 1996). A structural comparison of these two complexes suggests that gp40 and NKG2D receptor compete for binding to RAE-1γ as suggested earlier (Wang et al., 2012; Zhi et al., 2010). To investigate whether this is the case, we transfected wild type HEK293T with HA-RAE-1γ together with gp40WT or gp40LM and stained the cells with anti-HA antibody or with a recombinant mouse NKG2D-IgFc fusion protein. As expected, cells expressing gp40WT showed no RAE-1γ at the cell surface with both anti-HA and NKG2D stains. Strikingly, in the gp40LM-expressing cells, HA-RAE-1γ was present on the cell surface as detected by anti-HA stain, but a part of it was not recognized by the NKG2D stain (Fig. 5C), suggesting that gp40 interfered with its recognition.

To study whether this lack of RAE-1γ detection by NKG2D compromises NK cell activation, we used the previously established B78H1 cell lines expressing HA-RAE-1γ alone or with gp40WT or gp40LM. We subjected them to ^51^Cr release assays with freshly isolated murine NK cells. NK cell activation and specific lysis was the highest for B78H1 cells without gp40. Specific lysis of cells expressing either gp40WT or gp40LM was reduced, and it was about 20% less than cells expressing HA-RAE-1γ alone (Fig. 5D).

Our results suggest that the complex of gp40 and RAE-1γ is present at the cell surface, but RAE-1γ is masked by gp40 to abolish binding to the NKG2D receptor and thus limit NK cell activation.

## Discussion

### Retention of RAE-1γ parallels gp40-mediated MHC class I retention

NKG2D-mediated NK cell activation plays a crucial role in the defense against infection. The NKG2D receptor is expressed not only on NK cells but also on NK1.1^+^ T cells, γδ T cells, activated CD8^+^ αβ T cells, and activated macrophages (Diefenbach et al., 2000; Lanier, 2015). Multiple immunoevasins from both HCMV and MCMV suppress NKG2D activation by retaining NKG2D ligands inside the infected cell or by re-routing them for degradation. For most of these immunoevasins, the detailed mechanism of action remains elusive (Arapović et al., 2009b; Ashiru et al., 2009; Chalupny et al., 2006; Cosman et al., 2001; Eagle et al., 2009; Fielding et al., 2014, 2017; Hasan et al., 2005; Krmpotic et al., 2005; Lenac et al., 2006; Rölle et al., 2003; Wang et al., 2012; Wu et al., 2003; Zhi et al., 2010).

Our study, for the first time, explains at the molecular level how the retention of the NKG2D ligand, RAE-1γ, is achieved by MCMV gp40. We show, in accordance with the literature, that gp40 downregulates cell surface RAE-1γ by retaining it in the early secretory pathway (Fig. 1). We find that RAE-1γ retention depends on the interaction of gp40 with the host protein TMED10, in a way similar to MHC class I retention (Ramnarayan et al., 2018). If binding between TMED10 and gp40 is precluded (by using ΔTMED10 cells or by using gp40LM, a gp40 mutant that does not bind to TMED10), then RAE-1γ is no longer retained (Fig. 2–3).

### gp40 and RAE-1γ co-migrate through the early secretory pathway

For the first time, we show by co-immunoprecipitation that RAE-1γ and gp40 interact *in vivo*, which is consistent with the *in vitro* binding between both proteins that was reported earlier (Wang et al., 2012; Zhi et al., 2010). The complex is surprisingly strong, as it persists in 1% Triton X-100 (Fig 4). In contrast, gp40 and MHC class I molecules co-immmunoprecipitate only in milder conditions (1% digitonin), and the complex is sensitive to Triton X-100 (Janßen et al., 2016). This suggests that gp40 has a higher affinity to RAE-1γ than to MHC class I. Pulse-chase and co-immunoprecipitation analysis show that the gp40/RAE-1γ complex is formed immediately after protein synthesis and persists for at least two hours (Fig. 4).

If gp40 is not retained by TMED10, such as in TMED10 knockout cells or when gp40LM is used, then gp40 and RAE-1γ still bind to each other and migrate to the cell surface (Fig. 4) The final destination of the complex is, most likely, the lysosomes, as observed for gp40 by Ziegler et al., 2000.

The p24 family, of which TMED10 is a member, is involved in the ER export of some GPI-anchored proteins (Kaiser, 2000; Pastor-Cantizano et al., 2016). It is important to state that we found no evidence of direct binding between RAE-1γ, which is GPI-anchored, and TMED10 (Fig. 2).

In addition to RAE-1γ, gp40 also interacts with, and suppresses the function of, MHC class I and STING, but it is unclear how during infection, gp40 divides itself between its three target proteins (Arapović et al., 2009b; Janßen et al., 2016; Lodoen et al., 2003; Ramnarayan et al., 2018; Stempel et al., 2019; Wang et al., 2012; Zhi et al., 2010; Ziegler et al., 1997, 2000).

### Causes of gp40 surface expression

We and others have shown that the majority of gp40 is located in the early secretory pathway (Arapović et al., 2009b; Janßen et al., 2016; Ramnarayan et al., 2018; Stempel et al., 2019; Ziegler et al., 1997, 2000). The cell surface appearance of gp40 in viral infection (Fig. 4), with or without RAE-1γ bound to it, might have several reasons. First, gp40 that is produced in excess of available retention factors (i.e., p24 proteins) might travel to the cell surface by default. Second, gp40 molecules might be bound to TMED10, but TMED10 itself might lose its anchoring in the ER and travel to the surface in complex with gp40. We believe that the latter is possible since CMV remodels the secretory pathway to create the virus assembly compartment, to which some ER-resident proteins are relocated (Alwine, 2012; Procter et al., 2018; Tandon and Mocarski, 2012). In our work, we failed to detect TMED10 at the cell surface of MCMV-infected cells (not shown), but it was previously shown at the cell surface already 24 hours post infection, while its overall levels in the cell remained the same. The same was found for TMED9, another binding partner of gp40 (Nightingale et al., 2018; Ramnarayan et al., 2018). In the absence of CMV infection, TMED10 is mostly located in the early secretory pathway, involved in COPI- and COPII-dependent transport, but some studies have shown a fraction of TMED10 on the cell surface (Blum and Lepier, 2008; Gommel et al., 1999; Pastor-Cantizano et al., 2016; Zavodszky and Hegde, 2019).

### gp40 masks RAE-1γ at the cell surface

Our previous studies of MHC class I retention by gp40 show that if gp40 is not retained by TMED10, then MHC class I molecule surface expression is restored (Ramnarayan et al., 2018). In analogous conditions (*i.e*., no binding between gp40 and TMED10), we also detected cell surface HA-RAE-1γ with an anti-HA antibody. Surprisingly, though, this surface HA-RAE-1γ failed to stain with the monoclonal antibody, CX1. We attributed this lack of detectability by CX1 to a masking of the CX1 epitope of RAE-1γ by gp40 at the cell surface. The masking hypothesis is supported by the fact that both proteins, gp40 and RAE-1γ, are detected on the cell surface, and by the co-crystal structure of the gp40/RAE-1γ complex, in which gp40 binds to the top portion of the RAE-1γ molecule. We speculate that both proteins are present as a complex at the cell surface because they bind to each other tightly and co-migrate throughout the secretory pathway as a complex. In wild type cells expressing gp40LM, the CX1 epitope on RAE-1γ is also inaccessible, confirming that the cell surface RAE-1γ population was associated with gp40 (either WT or LM) that was preventing access to the CX1 antibody (Fig. 3–4).

### Masking of RAE-1γ prevents binding to NKG2D and NK cell activation

In the crystal structures of the gp40/RAE-1γ complex, gp40 is seen to cover the top (membrane-distal) surface of RAE-1γ (Wang et al., 2012). No structure of the complex of RAE-1γ with NKG2D exists, but there is a structure of RAE-1β, which shares 92% amino acid similarity with RAE-1γ, with NKG2D (Li et al., n.d.). In this complex, NKG2D binds to the same portion of RAE-1β that is covered by gp40 on RAE-1γ, suggesting that binding of NKG2D and gp40 to RAE-1γ is mutually exclusive (Wang et al., 2012). In agreement with this hypothesis, we observe a lack of recombinant NKG2D binding to cell surface RAE-1γ when gp40LM is present, and reduced ability of NK cells to recognize gp40-masked RAE-1γ and to kill the target cells (Fig 5). Thus, masking of surface RAE-1γ contributes to the impact of gp40 on NKG2D-dependent NK cell recognition of infected cells, and thus MCMV virulence (Lodoen et al., 2003).

Our results confirm the hypothesis of the group of Margulies, who suggested that gp40 might change the NKG2D binding site of RAE-1γ and/or mask RAE-1γ to prevent its recognition by NKG2D (Zhi et al., 2010). We propose that cell surface masking works as a backup mechanism to inhibit NK cell activation during infection (Fig. 6).

**Figure 6.**
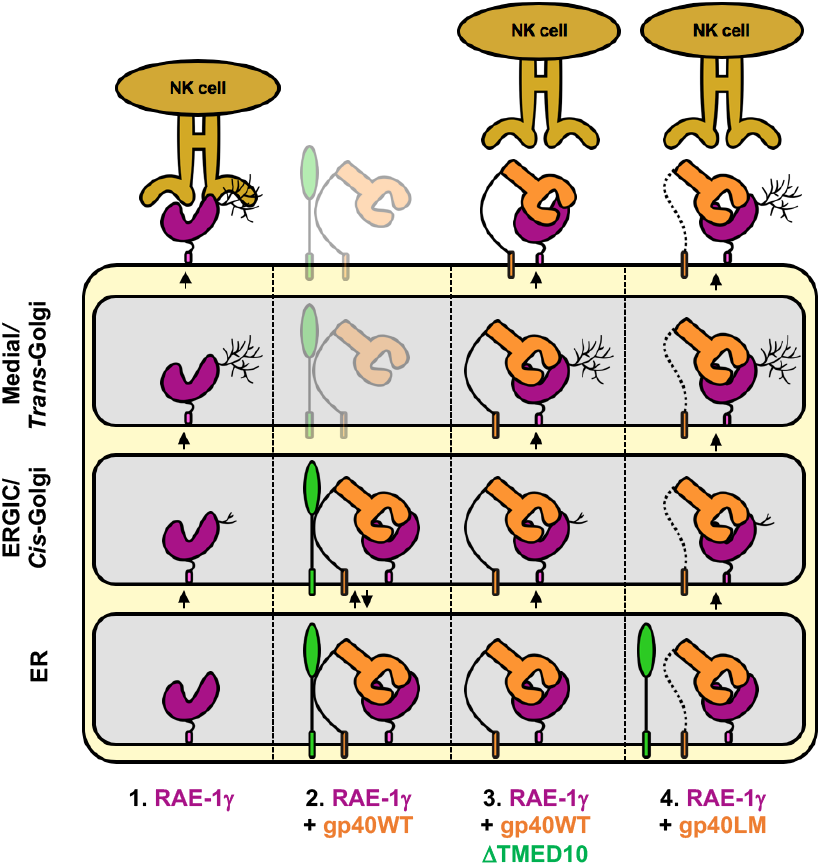
Proposed mechanism for gp40-mediated retention of RAE-1γ. 1. RAE-1γ expression without gp40. After synthesis, RAE-1γ (purple) is exported through the secretory pathway. During RAE-1γ transport, glycans are remodeled, making the glycosylation sensitive to EndoF1 in the ER, ERGIC, and *c/s*-Golgi, or resistant to EndoF1 beyond the medial Golgi. When RAE-1γ reaches the cell surface, it binds to the NKG2D receptor on the surface of the NK cell. 2. RAE-1γ expression in the presence of gp40WT. gp40 (orange) binds to RAE-1γ immediately after synthesis in the ER. The details of gp40/RAE-1γ binding are described in detail by Wang et al., 2012. Simultaneously, gp40 binds to TMED10 (green), a host molecule that carries ER retention/retrieval signals. In Ramnarayan et al., 2018 it was shown that the first three quarters of the 43 amino acid long gp40 linker mediate the binding, and we proposed that the gp40 linker binds to membrane proximal domain of TMED10. This model is yet to be validated. The trimeric complex of gp40, RAE-1γ and TMED10 circulates in the early secretory pathway (Ramnarayan et al., 2018). A small fraction of gp40 escapes the early secretory pathway and reaches the cell surface. It was shown by others that a fraction of TMED10 also reaches the cell surface (Nightingale et al., 2018). We do not know if both proteins might travel to the cell surface as a complex, or separately. 3. RAE-1γ expression in the presence of gp40WT, in the absence (knock out) of TMED10. gp40 binds to RAE-1γ immediately after synthesis. Both proteins are exported as a complex through the secretory pathway and reach the cell surface. RAE-1γ does not efficiently bind to the NKG2D receptor on NK cells because the interaction surface crucial for binding to the NKG2D is masked/covered by gp40. 4. RAE-1γ expression in the presence of gp40LM. The fate of gp40 and RAE-1γ is the same as in scenario 3. TMED10 is expressed in the cells but it cannot be bound by gp40, mimicking unavailability of TMED10 retention mechanism, without knocking out TMED10 itself.

One additional speculative mechanism is that gp40 travels to the cell surface on its own to bind to pre-existing RAE-1γ and inactivate it. This, however, seems unlikely because during viral infection, gp40 protein is detectable much earlier (3 hours post infection) than RAE-1 (18 hours post infection) (Tokuyama et al., 2011; Ziegler et al., 1997).

### Possible masking of other NKG2D ligands by immunoevasins

In addition to the masking of RAE-1γ by gp40 that we describe here, reports in the literature suggest that similar effects are possible for several other MCMV immunoevasins that prevent NKG2D receptor-mediated NK cell activation. This applies to MCMV m145, which regulates the NKG2D ligand MULT1 by interfering with its trafficking beyond the ERGIC/Golgi compartments; MCMV m155, which downregulates another NKG2D ligand, H60, by a proteasome-dependent mechanism; and MCMV *m138/fcr-1*, which interferes with the cell surface recycling of MULT1, H60, and RAE-1ε (the least gp40-susceptible RAE-1 isoform) and causes their degradation (Arapović et al., 2009a; Krmpotic et al., 2005; Lenac et al., 2006; Lodoen et al., 2004). Additionally, some HCMV immunoevasins that retain NKG2D ligands in the early secretory pathway by an unknown mechanism, namely UL16 and UL142, were detected at the cell surface (Ashiru et al., 2009; Vales-Gomez et al., 2006). All these immunoevasins may mask their targets on the cell surface in addition to their trafficking phenotype. This awaits further investigation.

Murine, and human, hosts of cytomegaloviruses have developed diverse NKG2D ligands that are induced upon viral infection. These ligands are an especially important target for CMV-mediated inactivation, and it seems likely that viruses would develop multiple mechanisms to inhibit NKG2D-mediated NK cell activation (Eagle and Trowsdale, 2007). It is conceivable that HCMV immunoevasins share these molecular mechanisms of inactivation, both the cell surface “masking” of host molecules and their intracellular retention by binding to p24 proteins.

A similar masking mechanism was observed for Ebolavirus spike glycoprotein (GP), a multifunctional protein responsible for host cell targeting, viral entry, and immune evasion. GP, which is heavily glycosylated, covers the surface proteins MHC class I and MICA (a human NKG2D ligand) to inhibit activation of T cells or NK cells, respectively. This shows that in addition to the downregulation of cell surface protein levels of the immune-activating host proteins, masking them on the cell surface is another immune evasion strategy shared by viruses from different taxonomic groups (Edri et al., 2018; Francica et al., 2010; Reynard et al., 2009).

## Materials and Methods

### Antibodies, reagents

Chemicals were purchased from AppliChem (Darmstadt, Germany) or Carl Roth (Karlsruhe, Germany). Mouse monoclonal hybridoma supernatants Y3 (Hammerling et al., 1982) and anti-HA 12CA5 (Niman et al., 1983) were as described previously. PE anti-mouse RAE-1γ antibody (CX1, 130107) was purchased from BioLegend (San Diego, USA). Anti-m152 (MCMV) (HR-MCMV-11) and Anti-m123/IE1 (MCMV) (HR-MCMV-12) were purchased from Capri (Canter for Proteomics, University of Rijeka, Croatia). Recombinant Mouse NKG2D-IgFc Chimera Protein, CF was purchased from R&D systems Germany (139-NK). Human IgG Fc APC-conjugated antibody (FAB110A) was purchased from R&D systems. Rabbit anti-calnexin serum was kindly provided by David Williams (Dept. of Biochemistry, University of Toronto, Toronto, Canada). Goat anti-Mouse IgG APC (115-135-164), Goat Fab anti-Rabbit IgG Alexa Fluor 488 (111-547-008) and Goat IgG anti-Rabbit IgG Cy3 (111-165-003) were purchased from DIANOVA GmbH Hamburg, Germany. FcR Blocking Reagent mouse (130-092-575) was purchased from Miltenyi Biotec GmbH Bergisch Gladbach, Germany. Protein Deglycosylation Mix II was purchased from New England Biolabs (P6044S).

### Cells

K41 cells (Gao et al., 2002) were kindly provided by Tim Elliott (Institute for Life Sciences, Southampton University Medical School); B78H1 murine melanoma cells deficient in MHC class I were a gift from Pier-Luigi Lollini (Curti et al., 2003) (Department of Specialized, Experimental, and Diagnostic Medicine, University of Bologna). Cells were grown at 37°C and 5% CO2 in high-glucose (4.5 g/L) DMEM (GE Healthcare) supplemented with 10% fetal calf serum (Biochrom, Berlin, Germany), 2 mM glutamine, 100 U/mL penicillin, and 100 mg/mL streptomycin.

### Retroviral Expression and Microscopy

Retroviral expression was performed as previously described (Hanenberg et al., 1996, 1997; Janßen et al., 2016; Hein et al., 2014). Immunofluorescence microscopy was performed as follows: cells were fixed with 4% para-formaldehyde and permeabilized with 0.1% Triton X-100. Next, the cells were stained and analyzed using laser scanning microscope. Antibody dilutions were as follows: goat anti-rabbit IgG Alexa Fluor 488 1:200, goat anti-rabbit IgG Cy3 1:200, rabbit anti-calnexin serum 1:300, CX1 1:200.

### Flow cytometry

For detection of cell surface proteins suitable antibodies were used: RAE-1γ, PE anti-mouse RAE-1γ antibody, CX1 or anti-HA 12CA5 or recombinant mouse NKG2D Fc Chimera Protein; gp40: anti-m152 antibody; MHC class I H-2K^b^, Y3. For detection of intracellular staining cells were fixed with 2.5% PFA for 20 min, permeabilized with 0.1% Triton X-100 for 15 min and stained with MCMV m123/IE1 antibody. The cells were harvested, stained, and then analysed using a CyFlow Space flow cytometer (Sysmex, Dresden, Germany). MCMV-infected cells were pre-treated with an FcR blocking reagent according to the manufacturer’s protocol before staining with antibodies. Antibody dilutions were as follows: Y3 and 12CA5 hybridoma supernatants 1:50, CX1 1:400, anti-m152 1:100, anti-m123/IE1 1:100, NKG2D-IgFc 1:20, secondary antibody APC 1:400.

### Pulse-chase experiments

Pulse-chase experiments were performed as previously described (Fritzsche and Springer, 2013). Briefly, cells were pulse-labeled with ^35^S labelling medium for 10 min and chased for the indicated times and lysed in 1% Triton X-100. HA-RAE-1γ was immunoprecipitated with anti-HA tag antibody (12CA5) and protein A agarose, digested with EndoF1 or New England Biolabs Protein Deglycosylation Mix II (removes all N-linked and O-linked glycosylation) as indicated, and analyzed using 12% SDS-PAGE and autoradiography. Gels were quantified using ImageJ (Wayne Rasband, NIH, USA).

### Co-immunoprecipitation and Re-immunoprecipitation

Labeling, pulse-chase, and immunoprecipitation (against the HA tag of HA-RAE-1γ) were performed as described before. Cells were lysed in buffer containing 1% digitonin or 1% Triton X-100. Precipitated proteins were eluted from the agarose beads by boiling in 50 μL denaturation buffer (1% SDS, 2 mM DTT) at 95°C for 10 min. Samples were cooled on ice, and SDS was neutralized with a 20-fold volume (1 mL) of 0.1% Triton X-100 in PBS. Samples were centrifuged at 1,000 x g for 10 min, and 900 μL was transferred to protein-A-agarose beads pre-bound with the antibody for re-immunoprecipitation and incubated for 1 hour at 4° C rotating. The beads were washed twice in PBS with 0.1% Triton X-100, and precipitated proteins were eluted by boiling in 20 μL denaturation buffer at 95°C for 10 min for SDS-PAGE followed by autoradiography.

### Viral infection

K41 cells were mixed with MCMV using MOI 0.5, subjected to centrifugal inoculation protocol at 2000 g for 30 min, infected for 3 hours, and incubated for indicated time. Infection was controlled using intracellular anti-m123/IE1 staining (dilution 1:100).

### NK cell generation

Single-cell suspension from spleens was depleted of erythrocytes, and NK cells were positively sorted using anti-DX5^+^ magnetic beads, according to the manufacturer’s instructions (Miltenyi Biotec, Bergisch Gladbach, Germany). Cells were resuspended in complete medium (RPMI; 10 mM HEPES, 2 × 10^-5^ M 2-ME, 10% FCS, 100 U/ml penicillin, 100 U/ml streptomycin) with with 1000 U/ml IL-2 (PeproTech) for 4 days.

### Cytotoxicity Assay

Target tumor cells were incubated for 1 h in the presence of Na_2_^51^CrO_4_ (Perkin Elmer, USA) and then washed thoroughly in PBS. NK cells and tumor cells were mixed at the ratios described, and after 4 h of coincubation, cell culture supernatants were taken and analyzed in a gamma-radiation counter (Wallac, Finland). Specific lysis was calculated according to the formula: percent specific lysis = [(experimental release–spontaneous release)/(maximum release– spontaneous release)] × 100.

### Statistical analysis

Data was analyzed by the use of GraphPad Prism 5.01 software (GraphPad, San Diego, California, USA). Levels of statistical significance were determined by Student T-test or one-way ANOVA, followed by Tukey post hoc tests. Values of p < 0.05 were considered statistically significant.

### Ethics statement

Mice on the C57BL/6 background were housed in isolated cages under specific pathogen free conditions at the Department of Microbiology, Tumor and Cell Biology and Astrid Fagraeus Laboratories, Karolinska Institutet, Stockholm. All procedures were performed under both institutional and national guidelines (Ethical numbers from Stockholm County Council N147/15).

## ACKNOWLEDGMENTS

We thank Linda Janssen for advice and help with the manuscript; Ursula Wellbrock for excellent technical assistance; and Miriam Herbert and Bersal Williams for additional laboratory work on this project.

## Author contributions

Conceptualization: N.L., S.S., R.V.R..; Methodology: N.L., S.S..; Validation: N.L.; Formal analysis: N.L.; Investigation: N.L., Z.H., S.G., B.C.; Resources: S.S., B.C., Writing: N.L., S.S., R.V.R.; Visualization: N.L.; Supervision: S.S.; Project administration: S.S.; Funding acquisition: S.S.

## List of Symbols and Abbreviations

HCMV: Human Cytomegalovirus
MCMV: murine cytomegalovirus
NK cell: natural killer cell
Endo F1: endoglycosidase F1
TMED10: transmembrane p24 trafficking protein 10
CX1: monoclonal anti-RAE 1γ antibody
gp40-: cells transfected with HA-RAE-1γ without gp40
gp40+: cells transfected with HA-RAE-1γ and gp40
gp40LM: gp40 linker mutant
gp40WT: gp40 wild type
PDM: Protein Deglycosylation Mix
APC: APC-conjugated secondary antibody
RE-IP: re-immunoprecipitation.

